# Comparative Proteomic Study of Retinal Ganglion Cells Undergoing Various Types of Cellular Stressors

**DOI:** 10.1101/2023.10.06.561236

**Authors:** Christopher R. Starr, James A. Mobley, Marina S. Gorbatyuk

## Abstract

Retinal ganglion cell (RGC) damage serves as a key indicator of various retinal degenerative diseases, including diabetic retinopathy (DR), glaucoma, retinal arterial and retinal vein occlusions, as well as inflammatory and traumatic optic neuropathies. Despite the growing body of data on the RGC proteomics associated with these conditions, there has been no dedicated study conducted to compare the molecular signaling pathways involved in the mechanism of neuronal cell death. Therefore, we launched the study using two different insults leading to RGC death: glutamate excitotoxicity and optic nerve crush (ONC). C57BL/6 mice were used for the study and underwent NMDA- and ONC-induced damage. Twenty-four hours after ONC and 1 hour after NMDA injection, we collected RGCs using CD90.2 coupled magnetic beads, prepared protein extracts, and employed LC-MS for the global proteomic analysis of RGCs. Statistically significant changes in proteins were analyzed to identify changes to cellular signaling resulting from the treatment. We identified unique and common alterations in protein profiles in RGCs undergoing different types of cellular stresses. Our study not only identified both unique and shared proteomic changes but also laid the groundwork for the future development of a therapeutic platform for testing gene candidates for DR and glaucoma.

## 1. Introduction

Retinal ganglion cells (RGCs) are neurons that play a crucial role in vision by transmitting visual information from the retina to the brain. There are several types of RGCs, each with a specific function in processing visual information. They receive input from photoreceptor cells, rods and cones, and integrate this information before sending it to the brain via the optic nerve. The axons of RGCs form the optic nerve, which carries visual signals to the brain, where signals are further processed to create visual perception. Once damaged, axons of the mammalian central nervous system (CNS) fail to regenerate. RGCs are of clinical relevance in diabetic retinopathy (Abu-El-Asrar et al., 2004), glaucoma (Joo et al., 1999), retinal arterial and retinal vein occlusions (Sucher et al., 1997) as well as optic neuropathies (Khan et al., 2021). In these diseases, RGCs may be irreversibly damaged. It is widely accepted that damaged RGCs not only serve as a model of optic neuropathies but also as the primary model for studying mechanisms of CNS axon degeneration and how therapies may promote regeneration.

After insult, both intrinsic and extrinsic factors contribute to whether CNS neurons survive, retain, or regenerate their axons. Although lowering intraocular pressure (IOP) is an approved treatment for glaucoma, it is not effective for every patient, and many other diseases affecting CNS neurons lack approved treatment options. In glaucoma, it has been reported that RGCs undergo apoptosis triggered by various factors, including autophagy, glutamate neurotoxicity, oxidative stress, neuroinflammation, immunity, and vasoconstriction (Kuehn et al., 2005; Shen et al., 2023). Autophagy can be induced by retinal hypoxia and axonal damage (Shen et al., 2023) while glutamate neurotoxicity is induced by the overstimulation of N-methyl-D-aspartate (NMDA) membrane receptors by glutamate, leading to progressive glaucomatous optic neuropathy (Shen et al., 2023).

In diabetes, retinal neuropathy involves progressive RGC death, axonal degeneration, and consequently, optic nerve degeneration (Bikbova et al., 2014). RGC loss occurs in diabetic patients even before the diagnosis of diabetic retinopathy. Furthermore, thinning of both the nerve fiber and the RGC layer has been documented in patients with diabetic retinopathy (Verbraak, 2014; Verma et al., 2009; Vujosevic and Midena, 2013) and animal diabetic models (Oleg S. Gorbatyuk, 2021; Pitale et al., 2021). One of the earliest experimental observations in animal models of diabetic retinopathy (DR) was the impairment of retrograde axonal transport in RGCs (Potilinski et al., 2020). Interestingly, this impairment was found to be even greater in type 1 diabetes than in type 2 diabetes, possibly due to a magnitude of metabolic dysfunctions contributing to optic nerve atrophy (Zhang et al., 1998). Under high glucose conditions, there was an observed increase in glutamate release, leading to significant extracellular glutamate accumulation and subsequent neurotoxicity of RGCs, further contributing to their deterioration (Ma et al., 2010).

Various approaches have been employed to analyze cellular signaling involved in RGC death and axonal deterioration. For instance, research groups have conducted unbiased proteomic screens of total mouse retina lysates (Hollander et al., 2012; Kwong et al., 2021; Magharious et al., 2011; Zhu et al., 2022) and used fluorescent-assisted cell sorting (FACS)(Belin et al., 2015) to isolate RGCs following optic nerve crush (ONC), one of the most extensively studied models of CNS axonal injury. However, these studies either examined whole retina lysates or assessed sorted RGCs at a time point (3 days after injury) when a significant amount of RGC apoptosis has already begun (Goldenberg-Cohen et al., 2012). Another common injury model is NMDA-induced excitotoxicity, a model mimicking the glutamate-induced excitotoxicity associated with multiple neurodegenerative diseases such as amyotrophic lateral sclerosis (ALS), ischemic stroke, and traumatic brain injury. To date, the only study examining proteomics in the context of NMDA-induced excitotoxicity of RGCs investigated proteomic changes in whole retinal lysates 12 hours after NMDA injection, which is a time point at which a substantial amount of RGC apoptosis is already in progress (Suo et al., 2022).

Understanding how CNS neurons respond to injury is crucial as we strive to develop therapeutic strategies for promoting neuroprotection, axon survival, and regeneration. Particularly, the significance of such studies lies in the development of neuroprotective interventions. Therefore, we initiated a comparative proteomic study, in which we analyzed the protein profiles of RGCs subjected to different cellular stress stimuli. The importance of our study lies in the fact that we not only identified differentially expressed proteins involved in two distinct cell death mechanisms but also revealed common biological processes that were similarly altered by different cell stress stimuli.

## 2. Results

### 2.1. Nontargeted quantitative proteomics of retinal ganglion cells

In our study, we adopted an approach to analyze molecular alterations in deteriorating RGCs before rampant cell death ensues, even though the peaks of apoptosis occur at different time points. To investigate early proteomic changes in RGCs following ONC-or NMDA-induced injury, protein lysates from Thy1-magnetic bead-isolated RGCs underwent nano high-performance liquid chromatography/mass spectrometry (LC/MS). RGC enrichment was confirmed via western blot (WB) using antibodies against RGC proteins, RNA-binding protein with multiple splicing (Rbpms), Class III β-tubulin (Tubb3/TUJ1), and photoreceptor markers, phosphodiesterase 6β (Pde6β) and rhodopsin (Rho) (**Fig. 1A**). As expected, RGC markers Rbpms and Tubb3/TUJ1 were highly enriched in RGCs and notably reduced in cells not bound by Cd90.2-coupled magnetic beads, corresponding to the rest of the retinal cells (**Fig. 1A**). Furthermore, photoreceptor proteins Pde6β and rhodopsin were present in trace amounts only in cells bound by the beads but highly expressed in unbound retinal cells. RGC enrichment was further validated by quantitative reverse transcriptase polymerase chain reaction (qRT-PCR) with primers against CD90/Thy1, Rho, and Pde6b (**Fig. 1B**). While *Thy1* was significantly elevated in cells bound by the magnetic beads and reduced in unbound cells, *Rho* and *Pde6b* mRNA levels were high in unbound cells and significantly diminished in bound cells. Together, the WB and qRT-PCR results indicate reliable RGC enrichment was achieved in our study.

**Figure 1.**
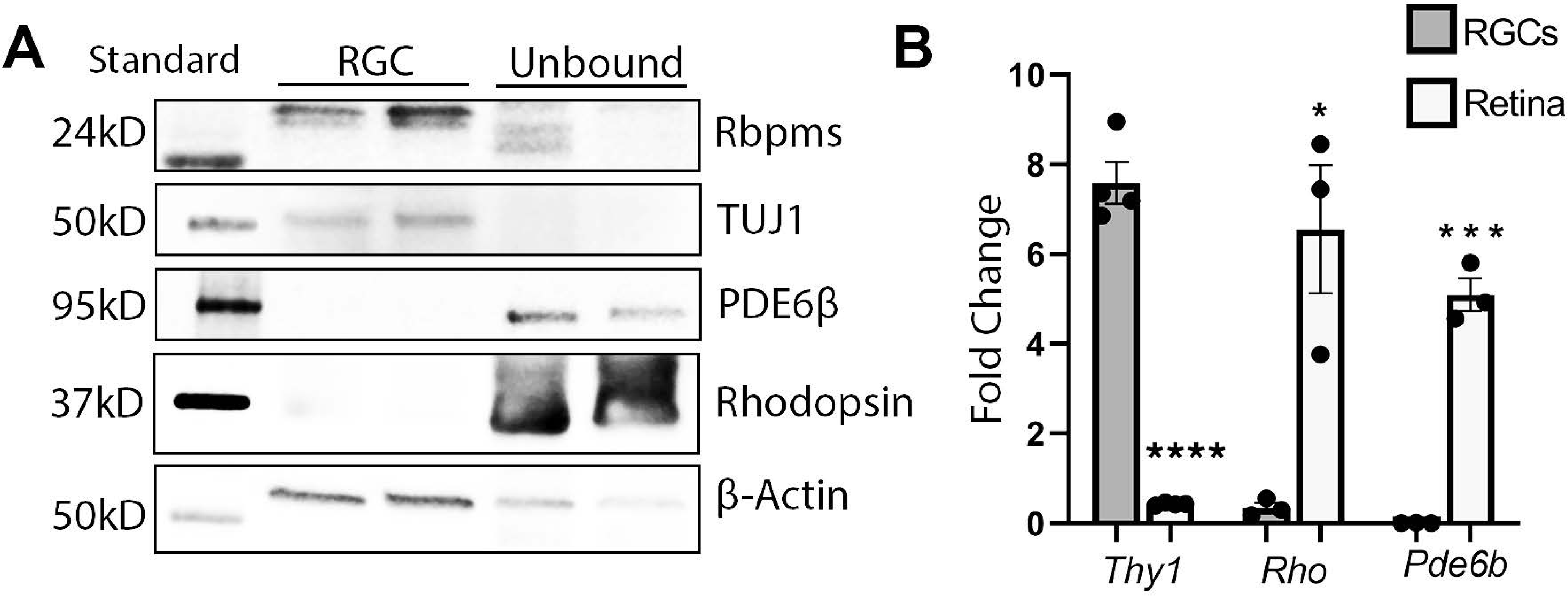
RGCs are enriched with Cd90.2 coupled magnetic beads. A) Western blot of select RGC and photoreceptor proteins indicating RGC enrichment. B) qRT-PCR analysis using primers against RGC or photoreceptor targets. * = p<0.05, *** = p<0.005, **** = p<0.001 (n=3-4). Data are shown as a standard deviation.

We analyzed identified proteins and prior to pathway analysis, grouped them according to the following parameters: 1) elevated in NMDA treated 2) reduced in NMDA treated 3) elevated in ONC treated 4) reduced in ONC 5) altered in NMDA and ONC.

### 2.2. NMDA treatment shifts RGC metabolic signaling

To induce excitotoxic damage to the retina, we intravitreally injected 1 µl of 20 mM NMDA in PBS, following a protocol previously described (Guo et al., 2021). RGCs were isolated 1 hour after NMDA injection and subjected to LC/MS analysis. Levels of many proteins were significantly altered following NMDA injection (see **Figure 2** and **Table 1** for top hits).

**Figure 2.**
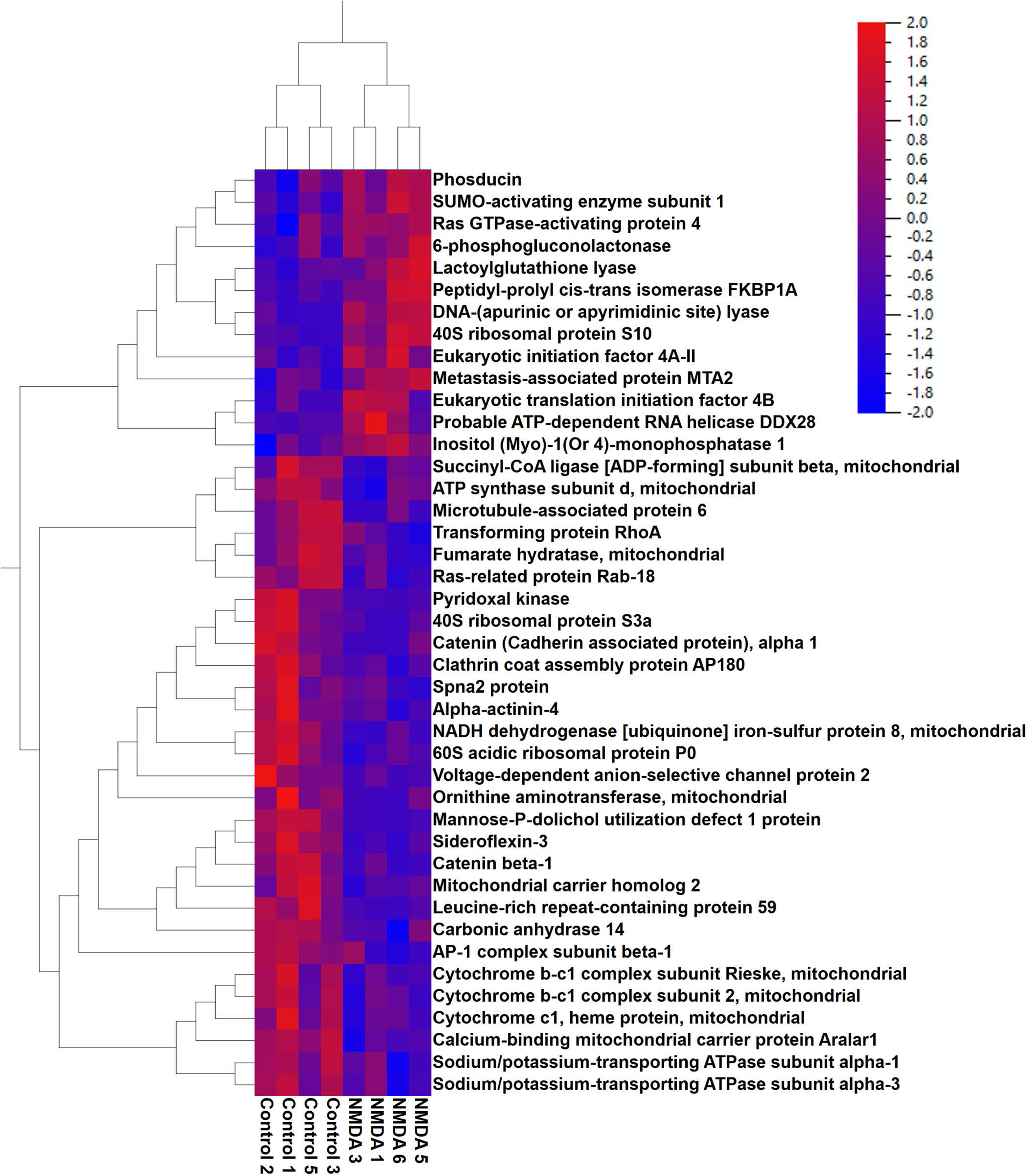
Proteins changed in RGCs 1 hour after following NMDA induced excitotoxicity. Heatmap depicting levels of protein top-hits.

**Table 1.** List of proteins most differentially expressed in RGCs 1 hour after NMDA treatment.

| **UniProtKB Names** | **Acc#** | **Fold(NMDA/C)** |
| --- | --- | --- |
| NADH dehydrogenase [ubiquinone] 1 alpha subcomplex subunit 12 | Q7TMF3 | 3.20 |
| 40S ribosomal protein S10 | P63325 | 3.20 |
| Peptidyl-prolyl cis-trans isomerase FKBP1A | P26883 | 2.93 |
| Selenium-binding protein 1 | P17563 | 2.68 |
| SUMO-activating enzyme subunit 1 | Q9R1T2 | 2.55 |
| Peptidyl-prolyl cis-trans isomerase FKBP4 | P30416 | 2.32 |
| Probable ATP-dependent RNA helicase DDX28 | Q9CWT6 | 2.28 |
| Translationally-controlled tumor protein | P63028 | 2.27 |
| Metastasis-associated protein MTA2 | Q9R190 | 2.24 |
| 60S ribosomal protein L35a | O55142 | 2.08 |
| Proteasome subunit alpha type-7 | Q9Z2U0 | 2.07 |
| 6-phosphogluconolactonase | Q9CQ60 | 2.06 |
| High mobility group protein HMG-I/HMG-Y | P17095 | 2.04 |
| Plasminogen activator inhibitor 1 RNA-binding protein | Q9CY58 | 2.03 |
| Phosducin | Q9QW08 | 1.98 |
| ATP-dependent RNA helicase DDX1 | Q91VR5 | 1.92 |
| Eukaryotic translation initiation factor 4B | Q8BGD9 | 1.90 |
| DnaJ homolog subfamily C member 8 | Q6NZB0 | 1.83 |
| ADP-ribosylation factor-like protein 3 | Q9WUL7 | 1.82 |
| Hypoxanthine-guanine phosphoribosyltransferase | P00493 | 1.81 |
| OTU domain-containing protein 7A | Q8R554 | 1.80 |
| Histone deacetylase complex subunit SAP18 | O55128 | 1.77 |
| Cytosolic non-specific dipeptidase | Q9D1A2 | 1.76 |
| High mobility group protein B3 | O54879 | 1.75 |
| Actin-related protein 2/3 complex subunit 3 | Q9JM76 | 1.73 |
| Inositol (Myo)-1(Or 4)-monophosphatase 1 | Q924B0 | 1.72 |
| DNA-(apurinic or apyrimidinic site) lyase | P28352 | 1.72 |
| Ras GTPase-activating protein 4 | Q6PFQ7 | 1.68 |
| Splicing factor 3A subunit 3 | Q9D554 | 1.67 |
| Lactoylglutathione lyase | Q9CPU0 | 1.62 |
| Cold-inducible RNA-binding protein | P60824 | 1.61 |
| Eukaryotic initiation factor 4A-II | E9Q561 | 1.59 |
| Ribosomal protein S23 | Q497E1 | 1.59 |
| Inositol polyphosphate 1-phosphatase | P49442 | 1.57 |
| Guanylate kinase | Q64520 | 1.55 |
| Prostaglandin E synthase 3 | Q9R0Q7 | 1.53 |
| Elongation factor 1-beta | O70251 | 1.53 |
| RNA-binding motif protein, X chromosome | Q9WV02 | 1.52 |
| Glutathione S-transferase P 1 | P19157 | 1.51 |
| Ribosomal protein S14 | O70569 | 1.50 |
| Sodium/potassium-transporting ATPase subunit alpha-3 | Q6PIC6 | -1.51 |
| Sodium/potassium-transporting ATPase subunit alpha-1 | Q8VDN2 | -1.54 |
| Glycogen phosphorylase, brain form | Q8CI94 | -1.55 |
| Hsc70-interacting protein | Q99L47 | -1.57 |
| N(G),N(G)-dimethylarginine dimethylaminohydrolase 1 | Q9CWS0 | -1.60 |
| Voltage-dependent anion-selective channel protein 2 | Q60930 | -1.61 |
| Cytochrome b-c1 complex subunit 2, mitochondrial | Q9DB77 | -1.62 |
| Protein NipSnap homolog 2 | O55126 | -1.63 |
| Very-long-chain 3-oxoacyl-CoA reductase | O70503 | -1.63 |
| Clathrin coat assembly protein AP180 | Q61548 | -1.63 |
| Mitochondrial carrier homolog 2 | Q791V5 | -1.64 |
| Thioredoxin-related transmembrane protein 1 | Q8VBT0 | -1.64 |
| ATP synthase subunit d, mitochondrial | Q9DCX2 | -1.64 |
| Leucine-rich repeat-containing protein 59 | Q922Q8 | -1.67 |
| Dihydrolipoyllysine-residue acetyltransferase component of pyruvate dehyd | Q8BMF4 | -1.67 |
| Cytoplasmic dynein 1 heavy chain 1 | Q9JHU4 | -1.68 |
| Synaptogyrin-3 | Q8R191 | -1.69 |
| Enoyl-CoA delta isomerase 1, mitochondrial | P42125 | -1.71 |
| MICOS complex subunit Mic60 | Q8CAQ8 | -1.71 |
| Ras-related protein Rab-18 | P35293 | -1.71 |
| Hexokinase-1 | P17710 | -1.74 |
| Sideroflexin-5 | Q925N0 | -1.75 |
| 60 kDa heat shock protein, mitochondrial | P63038 | -1.76 |
| Aldose reductase | P45376 | -1.77 |
| Fumarate hydratase, mitochondrial | P97807 | -1.79 |
| Transportin-1 | Q8BFY9 | -1.79 |
| Protein ERGIC-53 | Q9D0F3 | -1.79 |
| MAGUK p55 subfamily member 2 | Q9WV34 | -1.79 |
| Calcium-binding mitochondrial carrier protein Aralar1 | Q8BH59 | -1.80 |
| Ribophorin | Q61833 | -1.81 |
| Dynamin-3 | Q8BZ98 | -1.83 |
| Septin-7 | O55131 | -1.84 |
| F-actin-capping protein subunit alpha-1 | P47753 | -1.84 |
| Cytochrome c1, heme protein, mitochondrial | Q9D0M3 | -1.87 |
| Alpha-actinin-4 | P57780 | -1.90 |
| Spectrin beta chain, non-erythrocytic 1 | Q62261 | -1.90 |
| Succinyl-CoA ligase [ADP-forming] subunit beta, mitochondrial | Q9Z2I9 | -1.91 |
| Transforming protein RhoA | Q9QUI0 | -1.93 |
| Microtubule-associated protein 6 | Q7TSJ2 | -1.94 |
| Spna2 protein | B9EKJ1 | -1.94 |
| Isocitrate dehydrogenase [NAD] subunit, mitochondrial | Q91VA7 | -1.97 |
| Protein disulfide-isomerase A4 | P08003 | -1.97 |
| Protein NDRG2 | Q9QYG0 | -1.98 |
| Brain acid soluble protein 1 | Q91XV3 | -1.98 |
| Retinal dehydrogenase 1 | P24549 | -1.99 |
| Cleavage and polyadenylation-specificity factor subunit 6 | H3BJW3 | -1.99 |
| AP-1 complex subunit beta-1 | O35643 | -2.04 |
| AP-2 complex subunit alpha-2 | P17427 | -2.05 |
| Carbonic anhydrase 14 | Q9WVT6 | -2.06 |
| 40S ribosomal protein S3a | P97351 | -2.07 |
| S-formylglutathione hydrolase | Q9R0P3 | -2.07 |
| Pyruvate dehydrogenase E1 component subunit beta, mitochondrial | Q9D051 | -2.18 |
| 60S acidic ribosomal protein P0 | P14869 | -2.21 |
| Ornithine aminotransferase, mitochondrial | P29758 | -2.23 |
| Sideroflexin-3 | Q91V61 | -2.28 |
| Septin-5 | Q9Z2Q6 | -2.31 |
| CDGSH iron-sulfur domain-containing protein 1 | Q91WS0 | -2.33 |
| Sodium- and chloride-dependent GABA transporter 3 | P31650 | -2.35 |
| NADH dehydrogenase [ubiquinone] iron-sulfur protein 8, mitochondrial | Q8K3J1 | -2.38 |
| NADH dehydrogenase [ubiquinone] 1 alpha subcomplex subunit 6 | Q9CQZ5 | -2.47 |
| Dihydrolipoyl dehydrogenase, mitochondrial | O08749 | -2.48 |
| Synaptoporin | Q8BGN8 | -2.51 |
| Mannose-P-dolichol utilization defect 1 protein | Q9R0Q9 | -2.54 |
| Catenin (Cadherin associated protein), alpha 1 | Q6NV50 | -2.55 |
| Importin subunit beta-1 | P70168 | -2.59 |
| Phosphatidate cytidyltransferase 2 | Q99L43 | -2.60 |
| V-type proton ATPase 116 kDa subunit a isoform 1 | Q9Z1G4 | -2.61 |
| Cytochrome c, somatic | P62897 | -2.65 |
| Catenin beta-1 | Q02248 | -2.67 |
| Cytochrome b-c1 complex subunit Rieske, mitochondrial | Q9CR68 | -2.80 |
| AP-2 complex subunit beta | Q9DBG3 | -2.94 |
| Pyridoxal kinase | Q8K183 | -2.96 |
| Neuronal membrane glycoprotein M6-a | P35802 | -3.74 |

Proteins most significantly elevated following NMDA injection include NADH dehydrogenase [ubiquinone] 1 alpha subcomplex subunit 12, 40S ribosomal protein S10, Peptidyl-prolyl cis-trans isomerase FKBP1A, Selenium-binding protein 1, SUMO-activating enzyme subunit 1, Peptidyl-prolyl cis-trans isomerase FKBP4, Probable ATP-dependent RNA helicase DDX28, Translationally-controlled tumor protein, Metastasis-associated protein MTA2, 60S ribosomal protein L35a, and Proteasome subunit alpha type-7, among others. Proteins with the most significant reductions 1 hour after NMDA exposure include Neuronal membrane glycoprotein M6-a, Pyridoxal kinase, AP-2 complex subunit beta, Cytochrome b-c1 complex subunit Rieske-mitochondrial, Catenin beta-1, Cytochrome c-somatic, V-type proton ATPase 116 kDa subunit a isoform 1, Phosphatidate cytidylyltransferase 2, Importin subunit beta-1, Catenin (Cadherin associated protein)-alpha 1, and Mannose-P-dolichol utilization defect 1 protein (**Table 1)**.

The LC/MS results were then subjected to pathway analysis. NMDA treatment resulted in significant alterations in multiple pathways, with a notable impact on metabolic processes (**Table 2**). Pathways most reduced by NMDA injection were neutrophil degranulation, MHC class II antigen presentation, Sertoli cell junction signaling, oxidative phosphorylation, electron transport, TCA Cycle II, signaling by Retinoic Acid, COPI-mediated anterograde transport, and L1CAM interactions. Alterations in these pathways indicate a widespread shift in signaling in RGCs 1 hour after treatment with NMDA. Of note, many pathways involved in ATP production were reduced by NMDA mediated excitotoxicity. In support of these data, **Figure 3** provides a graphical overview of NMDA mediated mitochondrial dysfunction. Therefore, these findings demonstrate that NMDA treatment has a significant impact on metabolic signaling pathways in retinal ganglion cells, suggesting a link to impaired oxidative stress, and increased apoptosis.

**Figure 3.**
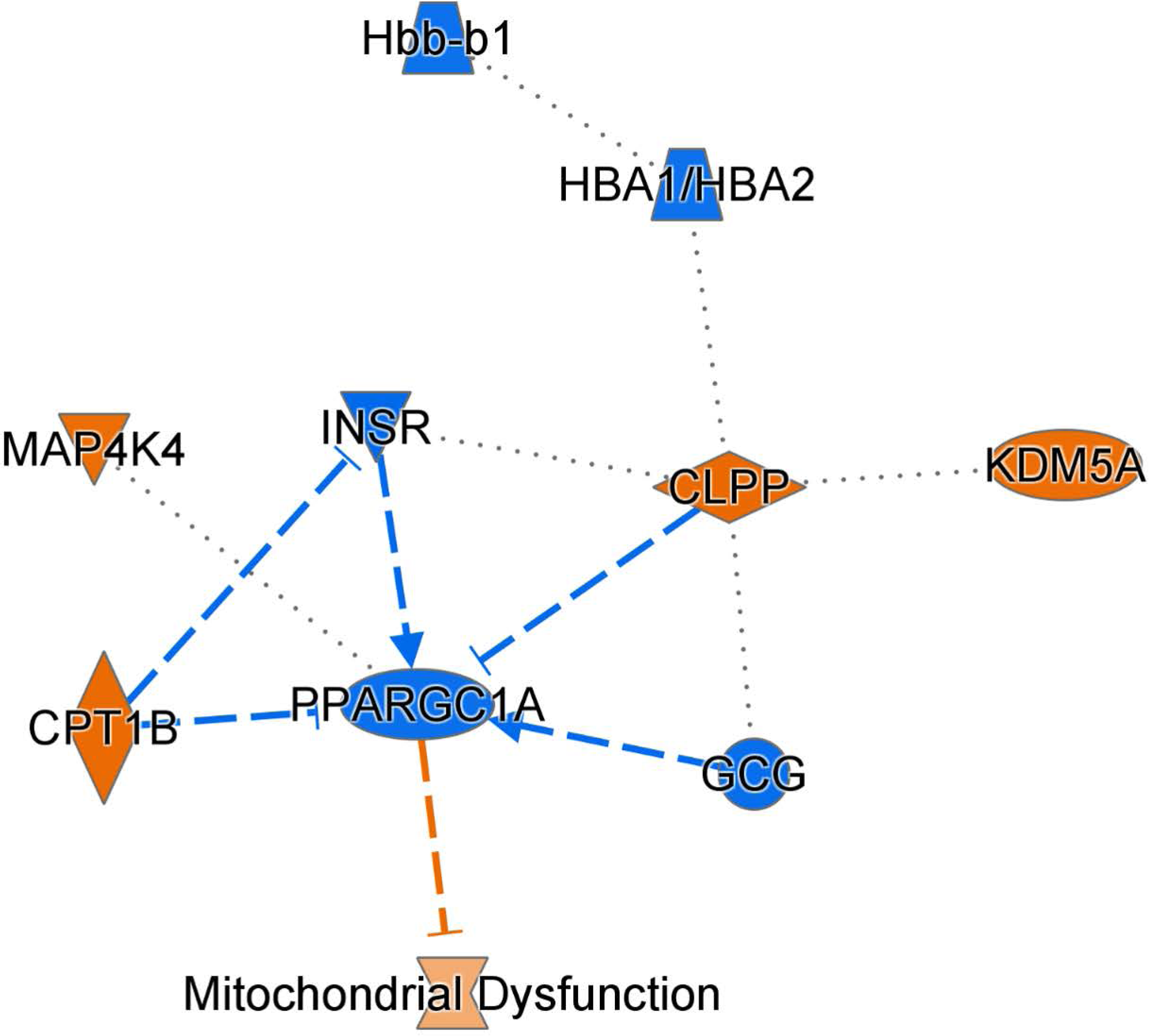
NMDA induced excitotoxicity induces alterations in metabolic signaling. Summary graph highlighting changes in mitochondrial signaling.

**Table 2.**
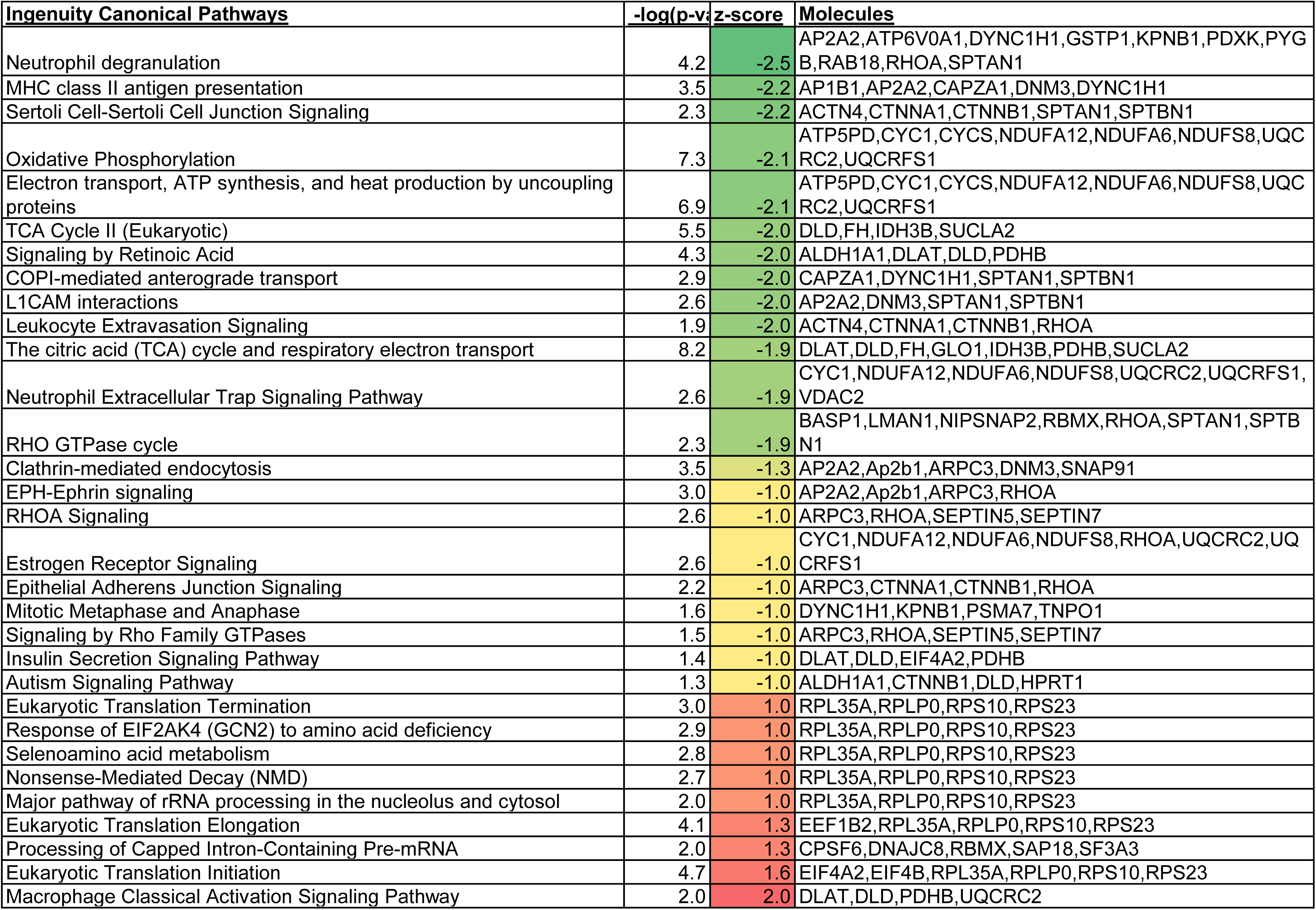
Ingenuity pathways significantly changed by NMDA-induced excitotoxicity.

Pathways that were most elevated by NMDA injection were Macrophage Classical Activation Signaling Pathway, Eukaryotic Translation Initiation, Processing of Capped Intron-Containing Pre-mRNA, Eukaryotic Translation Elongation, Major pathway of rRNA processing in the nucleolus and cytosol, Nonsense-Mediated Decay (NMD), Selenoamino acid metabolism, Response of EIF2AK4 (GCN2) to amino acid deficiency, and Eukaryotic Translation Termination **(Table 2)**. These results provide evidence about a shift in protein synthesis following excitotoxic injury (**Table 2**). Consistent with these findings, the levels of specific proteins related to these pathways, such as eukaryotic initiation factor (eIF) 4A2, eIF4B, and eukaryotic elongation factor (eEF) 1b2 were significantly elevated (**Table 1**). RNA processing perturbations are indicated by marked changes in multiple proteins including CPSf6, DNAJC8, RBMX. Collectively, our findings indicate that NMDA excitotoxicity in retinal ganglion cells leads to rapid perturbations in metabolic processes and mitochondrial homeostasis. These perturbations are characterized by a reduction in ATP production pathways. This shift reflects the intricate interplay between NMDA-induced damage and cellular metabolic responses, shedding light on the complex mechanisms involved in excitotoxicity-induced cellular alterations.

### 2.3. ONC results in metabolic dysfunction

C57BL/6 mice were subjected to ONC as described in a previously published protocol (Guo et al., 2021). RGCs were isolated 24 hours after ONC then subjected to LC/MS analysis. ONC led to alterations in several proteins and pathways (**Figure 4**, **Table 3**, and **Table 4**). The most elevated proteins were Splicing factor 3A subunit 1, CST complex subunit STN1, Stomatin-like protein 2-mitochondrial, DnaJ homolog subfamily C member 8, Arginine and glutamate-rich protein 1, Hepatoma-derived growth factor-related protein 2, Serine/arginine-rich splicing factor 6, Ddx3x protein, Probable ATP-dependent RNA helicase DDX28, RNA-binding protein FUS, and Proteasome subunit beta type-2 (Refer to Table 3 for complete list of top hits).

**Figure 4.**
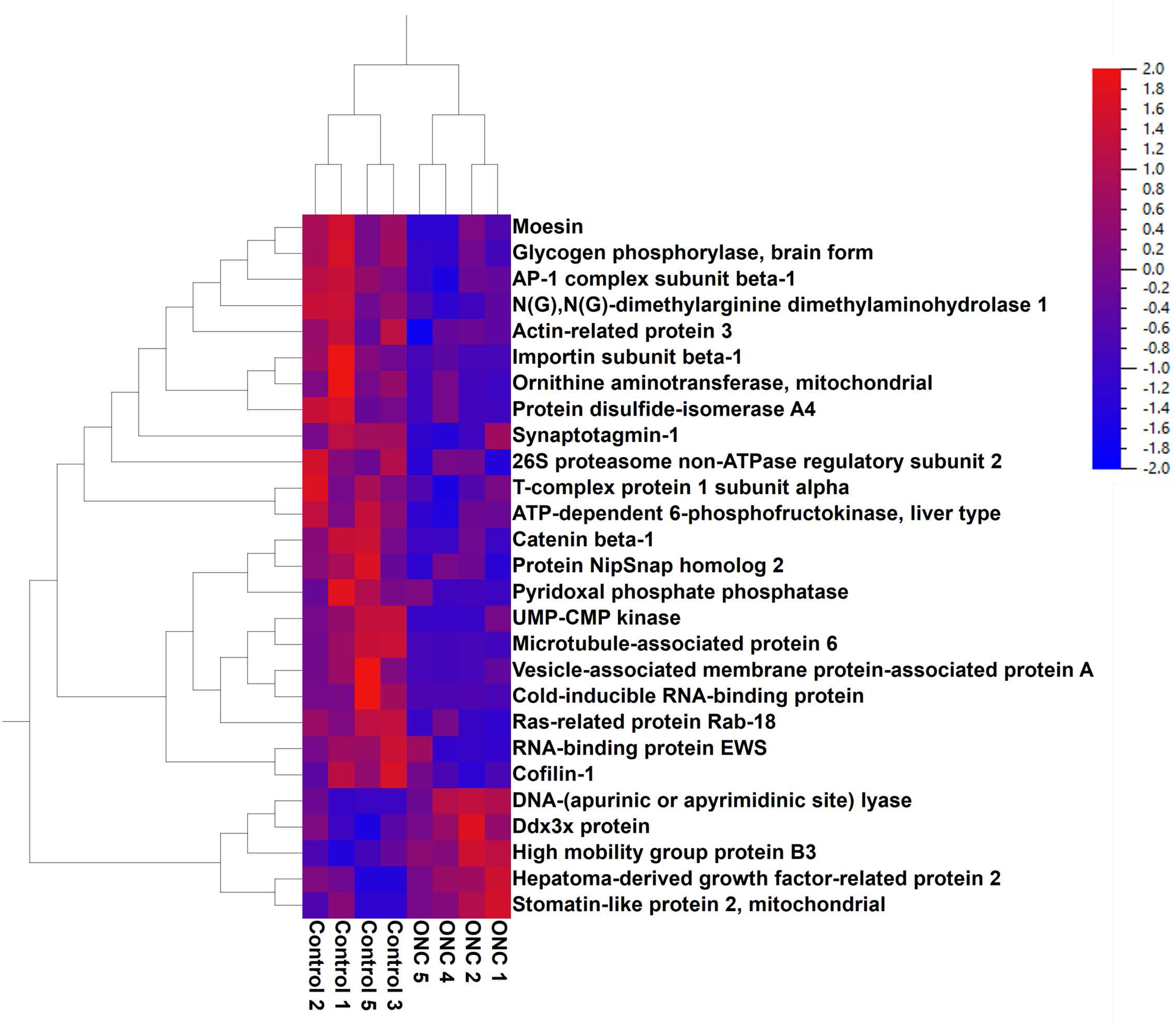
ONC results in changes in levels of proteins. Heatmap showing levels of proteins most changed 24 hours after ONC.

**Table 3.**
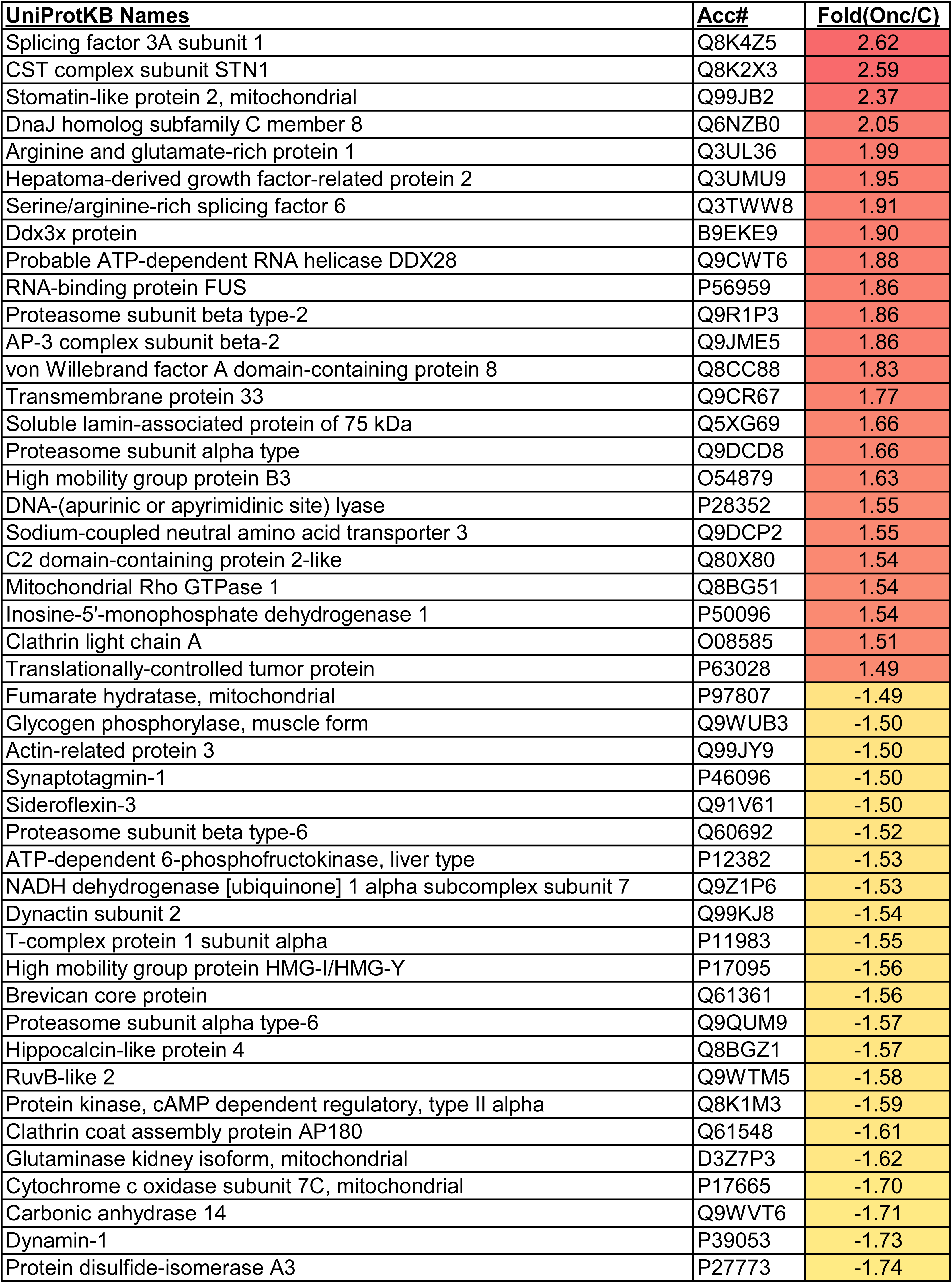
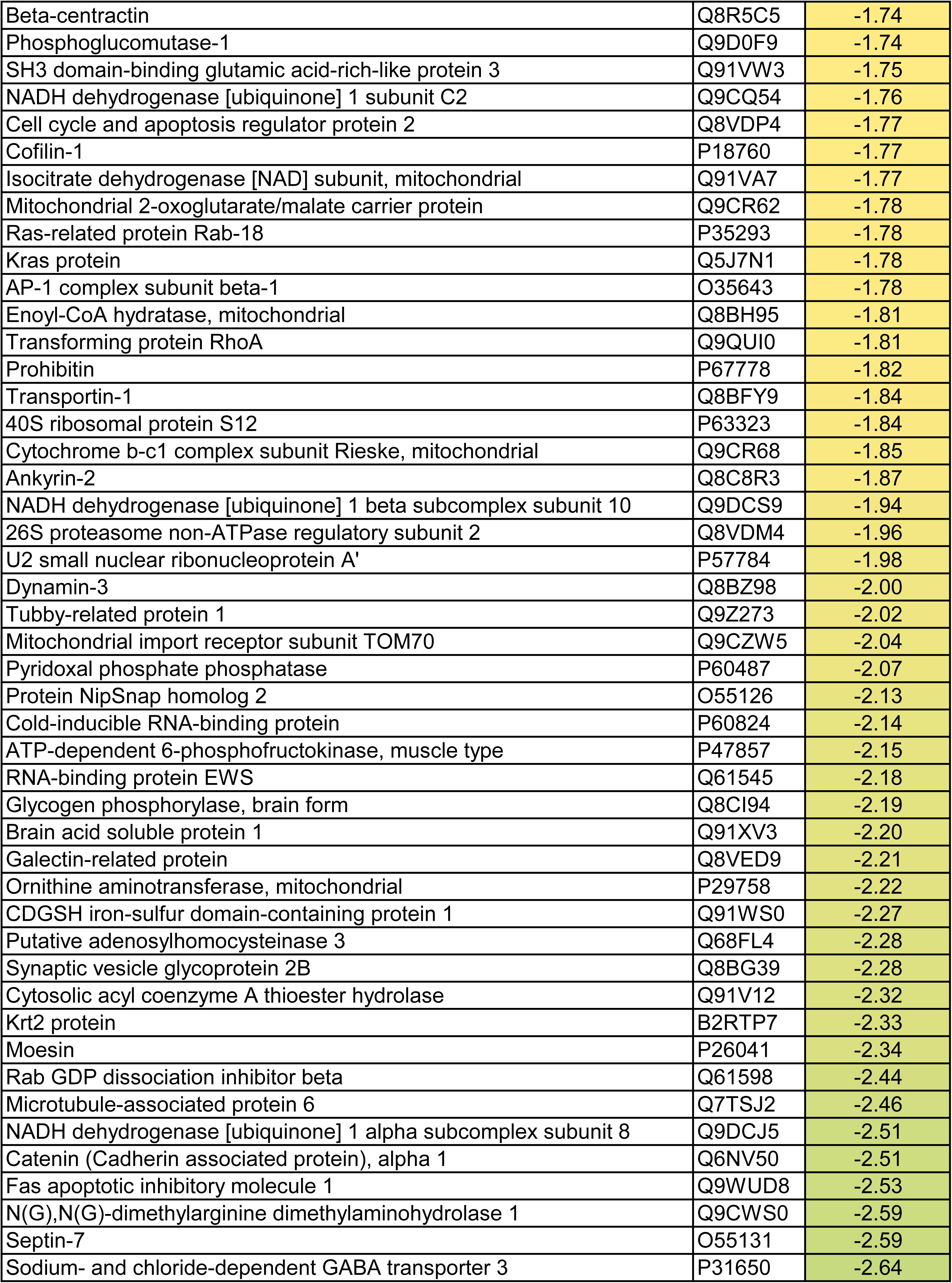
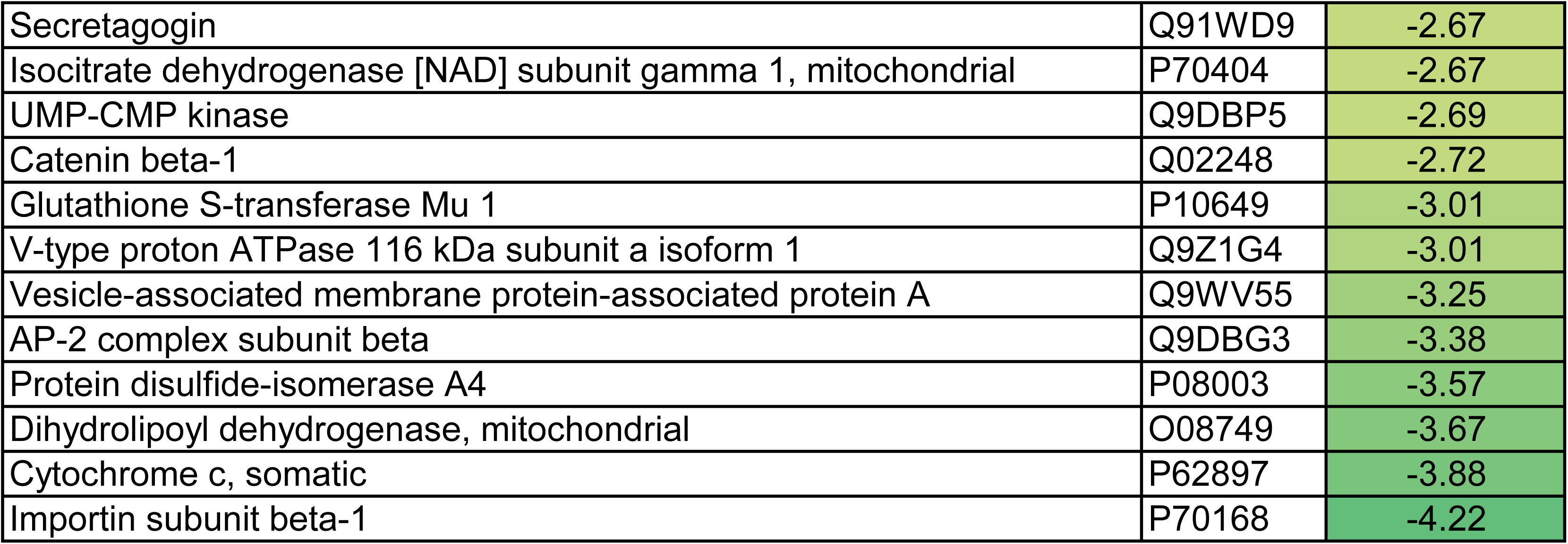
List of proteins with the largest fold-changes in RGCs 24 hours after ONC.

**Table 4.** List of ingenuity pathways significantly altered in RGCs 24 hours after ONC.

| Ingenuity Canonical Pathways | -log(p-value) | z-score | Molecules |
| --- | --- | --- | --- |
| Neutrophil degranulation | 7.8 | -2.7 | ACTR1B,ATP6V0A1,DDX3X,GDI2,IMPDH1,KPNB1,NDUFC2,PFKL,PGM1,P<br>SMD2,PYGB,RAB18,RHOA,VAPA |
| Electron transport, ATP synthesis, and heat production by uncoupling proteins | 4.7 | -2.4 | CYCS,NDUFA7,NDUFA8,NDUFB10,NDUFC2,UQCRFS1 |
| Eicosanoid Signaling | 2.9 | -2.4 | AP1B1,CTNNB1,KRAS,PDIA3,PRKAR2A,RHOA |
| Neutrophil Extracellular Trap Signaling Pathway | 2.1 | -2.4 | NDUFA7,NDUFA8,NDUFB10,PDIA3,TOMM70,UQCRFS1 |
| Oxidative Phosphorylation | 3.9 | -2.2 | CYCS,NDUFA7,NDUFA8,NDUFB10,UQCRFS1 |
| RHOA Signaling | 3.7 | -2.2 | ACTR3,CFL1,MSN,RHOA,SEPTIN7 |
| TCA Cycle II (Eukaryotic) | 5.6 | -2.0 | DLD,FH,IDH3B,IDH3G |
| The citric acid (TCA) cycle and respiratory electron transport | 4.0 | -2.0 | DLD,FH,IDH3B,IDH3G |
| Signaling by VEGF | 2.9 | -2.0 | CTNNA1,CTNNB1,KRAS,RHOA |
| RAC Signaling | 2.5 | -2.0 | ACTR3,CFL1,KRAS,RHOA |
| Opioid Signaling Pathway | 2.1 | -2.0 | Ap2b1,CLTA,CTNNB1,KRAS,PRKAR2A |
| Leukocyte Extravasation Signaling | 2.0 | -2.0 | CTNNA1,CTNNB1,MSN,RHOA |
| Ephrin Receptor Signaling | 1.9 | -2.0 | ACTR3,CFL1,KRAS,RHOA |
| Pancreatic Secretion Signaling Pathway | 1.7 | -2.0 | CA14,PDIA3,PRKAR2A,RHOA |
| Clathrin-mediated endocytosis | 5.8 | -1.9 | ACTR3,Ap2b1,CLTA,DNM1,DNM3,SNAP91,SYT1 |
| EPH-Ephrin signaling | 5.5 | -1.6 | ACTR3,Ap2b1,CFL1,CLTA,DNM1,RHOA |
| MHC class II antigen presentation | 4.8 | -1.6 | ACTR1B,AP1B1,CLTA,DCTN2,DNM1,DNM3 |
| Protein Kinase A Signaling | 2.7 | -1.6 | APEX1,CTNNB1,PDIA3,PRKAR2A,PYGB,PYGM,RHOA |
| Cardiac Hypertrophy Signaling (Enhanced) | 1.5 | -1.6 | APEX1,CTNNB1,KRAS,PDIA3,PRKAR2A,RHOA |
| Signaling by NTRK1 (TRKA) | 4.6 | -1.3 | CLTA,DNM1,DNM3,KRAS,RHOA |
| L1CAM interactions | 3.7 | -1.3 | Ank2,CLTA,DNM1,DNM3,MSN |
| Glioblastoma Multiforme Signaling | 3.1 | -1.3 | CTNNB1,KRAS,PDIA3,RHOA,RHOT1 |
| Signaling by Rho Family GTPases | 3.0 | -1.3 | ACTR3,CFL1,MSN,RHOA,RHOT1,SEPTIN7 |
| Actin Cytoskeleton Signaling | 2.4 | -1.3 | ACTR3,CFL1,KRAS,MSN,RHOA |
| Cardiac Hypertrophy Signaling | 2.3 | -1.3 | KRAS,PDIA3,PRKAR2A,RHOA,RHOT1 |
| Colorectal Cancer Metastasis Signaling | 2.2 | -1.3 | CTNNB1,KRAS,PRKAR2A,RHOA,RHOT1 |
| Serotonin Receptor Signaling | 1.3 | -1.3 | KRAS,PDIA3,PRKAR2A,RHOA,RHOT1 |
| Transcriptional regulation by RUNX3 | 6.7 | -1.1 | CTNNB1,KRAS,PSMA3,PSMA6,PSMB2,PSMB6,PSMD2 |
| Signaling by ROBO receptors | 6.2 | -1.1 | PRKAR2A,PSMA3,PSMA6,PSMB2,PSMB6,PSMD2,RHOA |
| Synaptogenesis Signaling Pathway | 4.2 | -1.1 | ACTR3,Ap2b1,CFL1,CTNNB1,KRAS,PRKAR2A,RHOA,SYT1 |
| Mitotic Metaphase and Anaphase | 4.1 | -1.1 | KPNB1,PSMA3,PSMA6,PSMB2,PSMB6,PSMD2,TNPO1 |
| RAF/MAP kinase cascade | 3.9 | -1.1 | KRAS,PHB1,PSMA3,PSMA6,PSMB2,PSMB6,PSMD2 |
| Chaperone Mediated Autophagy Signaling Pathway | 1.7 | -1.1 | ATP6V0A1,DDX3X,PSMA3,PSMA6,PSMB2,PSMB6,TPT1 |
| Gene and protein expression by JAK-STAT signaling after IL-12 stimulation | 4.7 | -1.0 | CFL1,MSN,SNRPA1,TCP1 |
| Actin Nucleation by ARP-WASP Complex | 3.2 | -1.0 | ACTR3,KRAS,RHOA,RHOT1 |
| Cargo recognition for clathrin-mediated endocytosis | 2.9 | -1.0 | Ap2b1,CLTA,SNAP91,SYT1 |
| ILK Signaling | 2.0 | -1.0 | CFL1,CTNNB1,RHOA,RHOT1 |
| Integrin Signaling | 1.9 | -1.0 | ACTR3,KRAS,RHOA,RHOT1 |
| Thrombin Signaling | 1.8 | -1.0 | KRAS,PDIA3,RHOA,RHOT1 |
| Docosahexaenoic Acid (DHA) Signaling | 1.6 | -1.0 | CYCS,PDIA3,PRKAR2A,SYT1 |
| Autism Signaling Pathway | 1.4 | -1.0 | CTNNB1,DLD,KRAS,PRKAR2A |
| Glutamatergic Receptor Signaling Pathway (Enhanced) | 1.3 | -1.0 | GLS,PDIA3,PRKAR2A,SLC38A3 |
| Gap Junction Signaling | 1.3 | -1.0 | CTNNB1,KRAS,PDIA3,PRKAR2A |
| Granzyme A Signaling | 3.5 | 1.0 | APEX1,NDUFA7,NDUFA8,NDUFB10 |
| Processing of Capped Intron-Containing Pre-mRNA | 1.5 | 1.0 | DNAJC8,SF3A1,SNRPA1,SRSF6 |
| Sirtuin Signaling Pathway | 5.3 | 1.6 | APEX1,GLS,NDUFA7,NDUFA8,NDUFB10,PFKM,SF3A1,TOMM70,UQCRFS1 |
| Mitochondrial Dysfunction | 4.7 | 1.7 | CYCS,DLD,NDUFA7,NDUFA8,NDUFB10,PRKAR2A,RHOT1,TOMM70,UQC<br>RFS1 |

Proteins most significantly reduced included Importin subunit beta-1, Cytochrome c-somatic, Dihydrolipoyl dehydrogenase-mitochondrial, Protein disulfide-isomerase A4, AP-2 complex subunit beta, Vesicle-associated membrane protein-associated protein A, V-type proton ATPase 116 kDa subunit a isoform 1, Glutathione S-transferase Mu 1, Catenin beta-1, and UMP-CMP kinase, to name a few (**Table 3**). Of note, ONC resulted in perturbations in metabolic signaling cascades. For example, electron transport, oxidative phosphorylation and citric acid cycle signaling were all reduced 24 hours after ONC while Macrophage Classical Activation Signaling pathway was elevated. Overall, the difference in the reduced cellular pathways in RGC with NDMA and ONC demonstrated a few similarities.

For example, the pathways most significantly reduced by ONC induced excitotoxicity were Neutrophil Degranulation, Electron transport, Eicosanoid Signaling, Neutrophil extracellular trap signaling, Oxidative phosphorylation, RHOA signaling, TCA Cycle II, The citric acid cycle and respiratory electron transport, signaling by VEGF, and RAC signaling, among others (**Table 4**).

Signaling most elevated following ONC injury was mitochondrial dysfunction, sirtuin signaling, Processing of Capped Intron-Containing Pre-mRNA, Granzyme A Signaling, Glutaminergic Receptor Signaling Pathway (Enhanced), Autism Signaling Pathway, Docosahexaenoic Acid (DHA) Signaling, Thrombin Signaling, and Integrin Signaling (**Table 4**). **Figure 5** shows the graphical summary of altered signaling pathways in the retina with ONC. As we can see, the major changes in RGC with ONL occurs in cytoskeletal organization.

**Figure 5.**
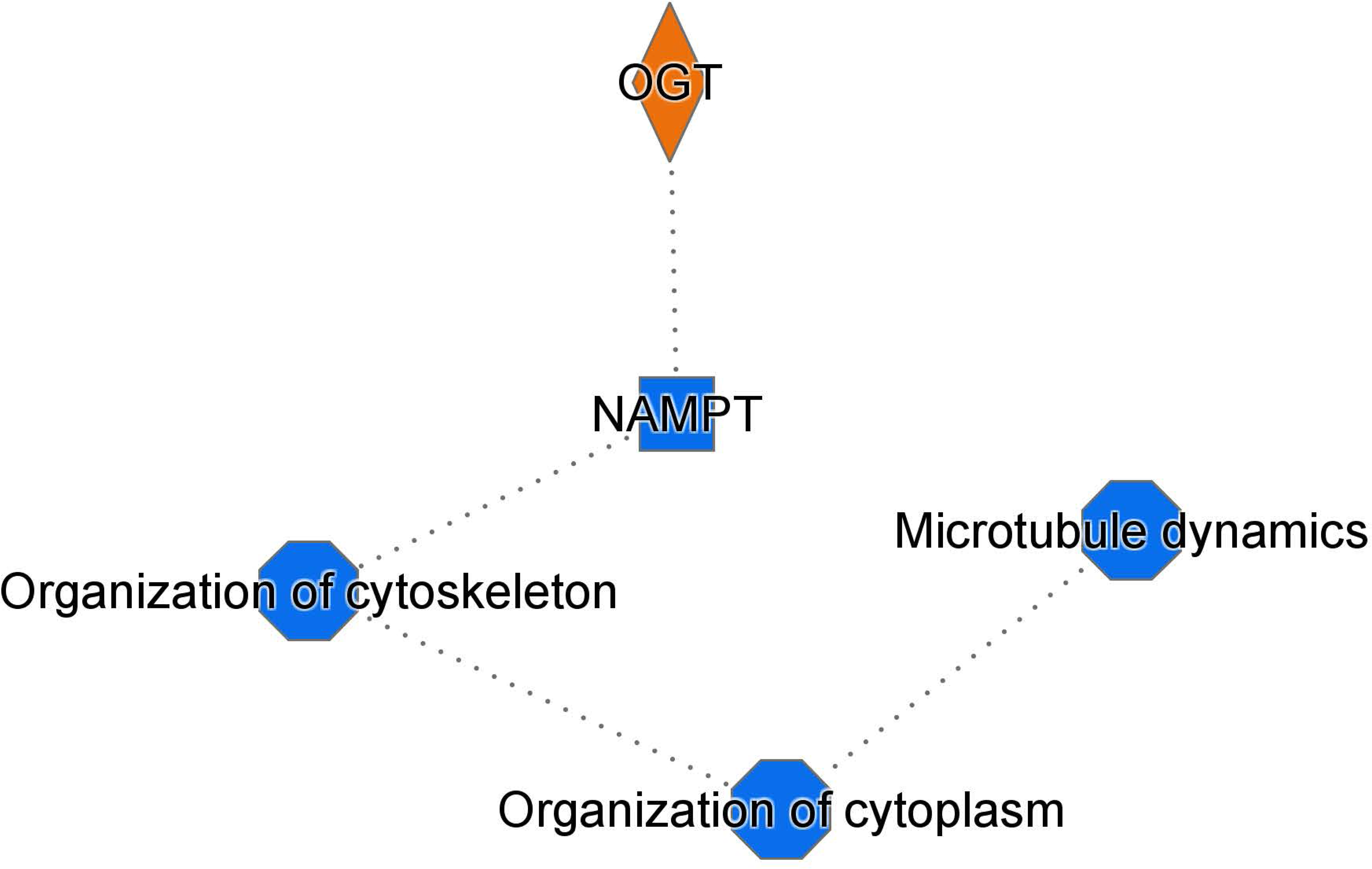
ONC leads to changes in cellular organization. Graphical summary detailing changes in cellular organization following ONC.

### 2.4. ONC and NMDA induced excitotoxicity both alter a subset of proteins

One of the primary objectives of this study was to identify proteins that exhibit changes in multiple models of neuronal injury. In pursuit of this goal, we successfully identified proteins that exhibited alterations in both models (**Fig. 6** and **Table 5**). Remarkably, the majority of these protein level changes, whether upregulation or downregulation, were notably consistent across both groups. Specifically, the most significantly downregulated proteins in both groups were importin subunit beta-1, Cycs, dihydrolipoyl dehydrogenase, protein disfulfide-isomerase a4 (Pdia4), AP-2 complex subunit beta, V-type proton ATPase 116 kDa subunit a isoform 1, Catenin beta-1, Sodium- and chloride-dependent GABA transporter 3, Septin-7, N(G),N(G)-dimethylarginine dimethylaminohydrolase 1, Catenin (Cadherin associated protein)-alpha 1, Microtubule-associated protein 6, and CDGSH iron-sulfur domain-containing protein 1. The most significantly downregulated proteins were alongside significant downregulation of DnaJ homolog subfamily C member 8 (Dnajc8), Probable ATP-dependent RNA helicase DDX28, High mobility group protein B3 (Hmgb3), DNA-(apurinic or apyrimidinic site) endonuclease (Apex1), and Translationally-controlled tumor protein (Tpt1) in both treated RGCs when compared to the control group. To validate the LC/MS proteomics, we assessed levels of Cytochrome C somatic (Cycs), a protein reduced in both ONC and NMDA groups, by western blot (Figure 6B and C). As expected, we also observed a significant reduction in Cycs by western blot. **Table 5**, which provides a visual representation of these modified protein levels, highlights the proteins significantly altered following injury. Two proteins exhibited an opposite trend in the two models. Both Cold-inducible RNA-binding protein and High mobility group protein HMG-I/HMG-Y were upregulated in NMDA challenged RGCs but downregulated following ONC.

**Figure 6.**
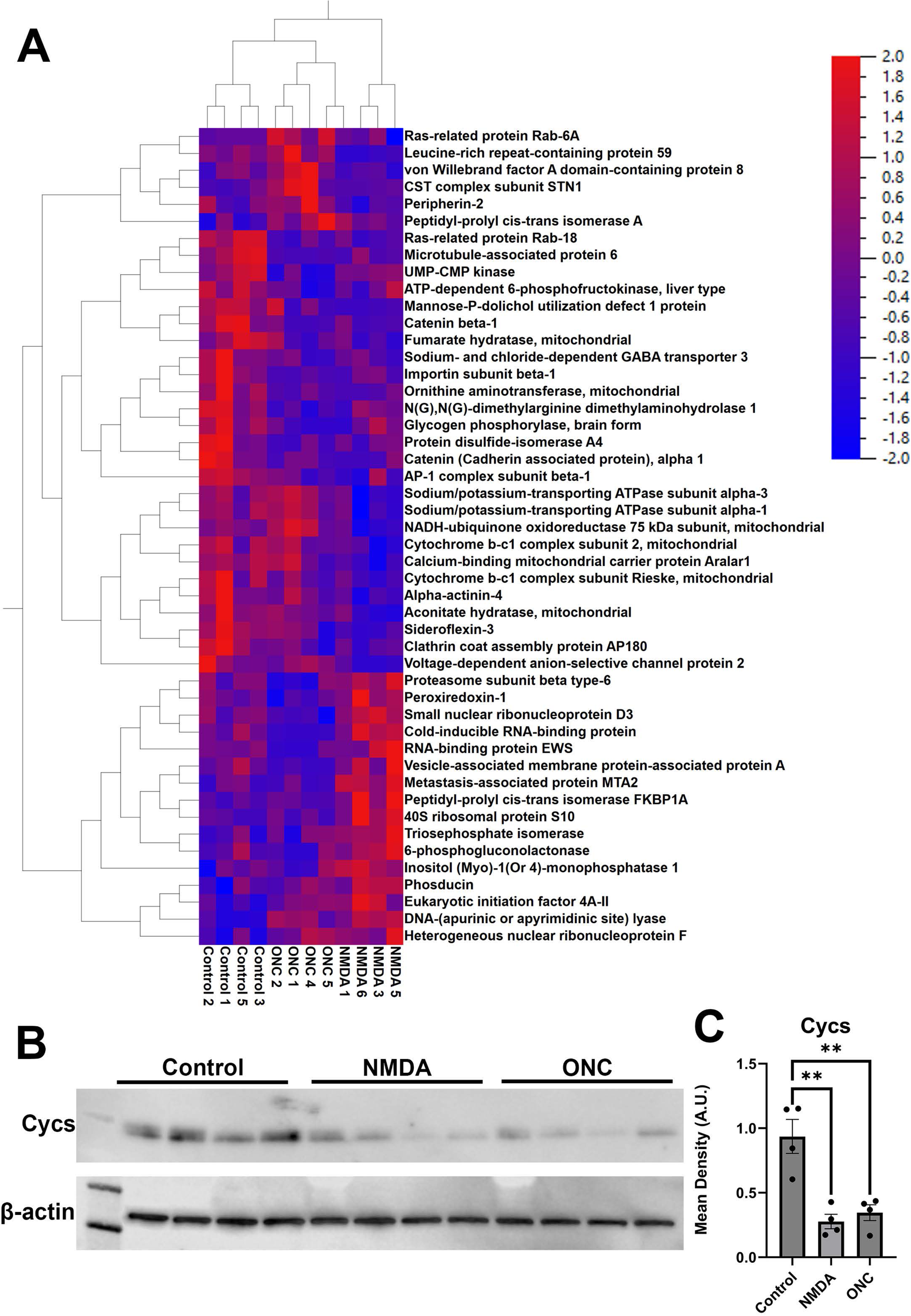
Proteins changed in RGCs 1 hour after NMDA induced excitoxicity as well as 24 hours after ONC. A) Heatmap identifying top-hits. B) Western Blots of protein lysates from enriched RGCs. C) Graph depicting Cycs levels normalized to β-actin. A.U. = arbitrary units. **=p<0.01

**Table 5.**
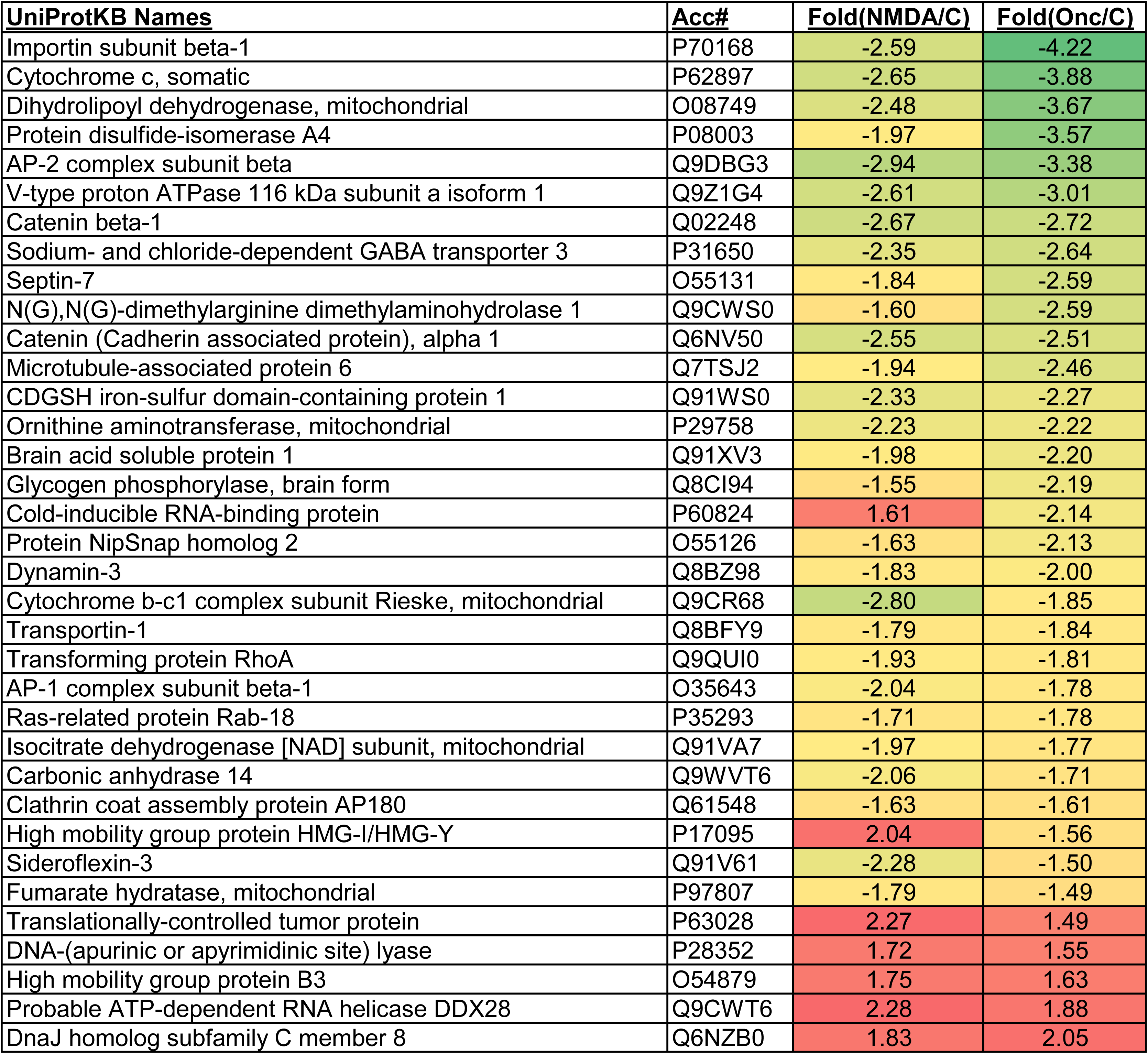
Proteins significantly changed in RGCs in both treatment groups, NMDA and ONC.

The analysis of differentially expressed proteins revealed 28 proteins with significantly reduced expression, 5 proteins with significantly increased expression, and two proteins exhibiting an opposite response pattern to the stimulus in both types of treated RGCs.

## 3. Discussion

In the present study, we examined acute changes in RGC global protein expression following NMDA injection or ONC. We provide evidence that NMDA excitotoxicity and ONC alters multiple signaling pathways, many of which are metabolic. We identified common proteins and pathways manifesting a decline and elevation in expression.

NMDA excitotoxicity is a model glutamate excitotoxicity. Glutamate excitotoxicity, along with glutamate dysregulation, the primary means by which RGCs are theorized to die in glaucoma and other retinal diseases including diabetic retinopathy (DR). For example, impairment of retrograde axonal transport and glutamate release have been found in animal models of DR (Potilinski et al., 2020). Classically, the primary cause of glutamate excitotoxicity is ATP depletion and impaired glutamate transport, resulting in a buildup of extracellular glutamate, leading to excitotoxicity by overloading NMDA receptor (NMDAR)-expressing cells with Na^+^ and Ca^2+^. Glutamate excitotoxicity is known to be involved in various diseases, including Alzheimer’s (Tannenberg et al., 2004), Parkinson’s (Verma et al., 2022), or Huntington’s (Girling et al., 2018), and predicted to be involved in choroidal vessel occlusion, glaucoma, and diabetic retinopathy (Lucas and Newhouse, 1957; Massey and Miller, 1987, 1990; Thoreson and Witkovsky, 1999; Yumnamcha et al., 2020). Elevated Ca^2+^ can lead to changes in several Ca^2+^-sensitive signaling cascades and eventually mitochondrial-mediated apoptosis (Verma et al., 2022). An increase in intracellular Na^+^ can lead to cell swelling. NMDA-mediated excitotoxicity leads to synaptic degeneration and dendritic pruning before cell death. By isolating RGCs one hour after NMDA injection, we hoped to see how the RGCs are altered in the time window before full-fledged apoptosis begins. We did indeed observe perturbations in several biological processes after NMDA injection. It is possible that the calcium elevation that occurs during excitotoxicity could be the major driving force behind excitotoxic cell death. Cells respond to this is by sequestering this excess calcium in the endoplasmic reticulum and mitochondria. Mitochondria is the site of oxidative phosphorylation, a process that generates ATP and is regulated by calcium (Glancy et al., 2013). This is consistent with our findings on a reduction of signaling cascades related to ATP production.

The murine optic nerve crush injury model is the most commonly used model of RGC injury that mimics molecular events occurring upon traumatic optic neuropathy, glaucoma, etc. (Tang et al., 2011). Moreover, the prevalence of optic neuropathy in diabetics could increase with diabetes duration (Hua et al., 2019). In this model, the crush injury to the optic nerve leads to retinal ganglion cell apoptosis. Like NMDA-induced excitotoxicity, this model can also be used to study the general processes and mechanisms of neuronal death and survival, which is essential for the development of therapeutic measures. In addition, pharmacological and molecular approaches can be used in this model to identify and test potential therapeutic reagents to treat different types of optic neuropathy. Interventions that are neuroprotective for RGCs challenged with ONC are usually neuroprotective for other CNS neurons (Le Pichon et al., 2017). In addition, axon regeneration promoting treatments discovered with the ONC model can typically promote axon regeneration in the spinal cord (Bhowmick and Abdul-Muneer, 2021). Following ONC, axonal components briefly travel retrogradely prior to the axons returning to the site of injury, only to fail to pass the crush site. In fact, knocking out an inhibitor of the retrograde injury response of RGCs, dual-leucine zipper kinase (DLK), leads to robust RGC survival (Watkins et al., 2013), indicating that retrograde signaling is crucial for cell death signaling following ONC. Based on this, in the earliest response to ONC, we would anticipate changes in the machinery responsible for transport and localization. As expected, we did identify reductions in GO biological processes pertaining to the localization of organelles cellular proteins and cellular macromolecules. After injuring the axon with ONC, we expected to observe changes in synaptic signaling. These changes are presented by the decline in multiple proteins including Catenin beta-1, Sodium- and chloride-dependent GABA transporter 3, AP180 and Synaptotagmin-1. Together, these changes suggest that, as expected, damaging the axon not only disrupts axonal components but also the cellular localization of various cellular entities.

In both RGC degenerative models, we identified common proteins that responded to cellular insults in a similar manner, as well as proteins that exhibited a unique decline in expression in both challenged RGCs. While our current study has its limitations due to the lack of validation of the 33 proteins similarly altered in both groups (**Table 5**), future investigations should prioritize the validation of the roles of these proteins in RGC survival. In particular, it would be interesting to assess the role of these proteins in RGC survival during the development of diabetic retinopathy or glaucoma. For example, Rab18 deficiency detected in both types of challenged RGCs is the molecular deficit underlying Warburg micro syndrome, characterized by eye, nervous system, and endocrine abnormalities (Handley and Sheridan, 1993). Moreover, a recent study highlighted the involvement of Rab18 in lipid metabolism in human diabetic adipose tissue and demonstrated that Rab-18 contributes to insulin resistance in obese individuals. (Guzman-Ruiz et al., 2020; Pulido et al., 2011) Another example is DnaJC8 protein, the effect of which is closely associated with the aggregation of polyQ-containing proteins in a cellular model of spinocerebellar ataxia type 3 (SCA3).(Ito et al., 2016) The authors have shown that DnaJC8 overexpression significantly reduces polyQ aggregation and apoptosis. Therefore, DnaJC8 should be validated for its neuroprotective role in the survival of RGCs undergoing both cellular stressors.

## 4. Conclusion

In summary, our study was designed to not only identify both individual and shared proteomic changes in retinal ganglion cells undergoing different stress stimuli just before initiating a pro-apoptotic cell death program but also to lay the groundwork for the future development of a therapeutic platform. Future studies should therefore concentrate on validating common proteome changes associated with RGC deterioration in both cell death models. This approach would enhance our understanding of the underlying mechanisms driving RGC degeneration, potentially uncovering key biomarkers or therapeutic targets crucial for developing effective treatments for conditions such as glaucoma, diabetic retinopathy, and optic neuropathies.

## 5. Methods

### 5.1. Animals

All animal procedures were approved by The University of Alabama at Birmingham institutional animal use and care (IACUC) committee and in accordance with the statement for the Use of Animals in Ophthalmic and Vision Research by The Association for Research in Vision and Ophthalmology (ARVO). This study is reported in accordance with ARRIVE guidelines. C57BL/6J mice were purchased from Jackson Laboratory (Bar Harbor, ME). An equal number of female and male mice were used in this study (10:10). No difference was observed between male and female mice. For all procedures, mice were anesthetized with ketamine (100 mg/kg) and xylazine (10mg/kg).

To induce injury to RGCs, mice were either intravitreally injected with 1µl of 20mM N-Methyl D aspartic acid (NMDA) or underwent optic nerve crush (ONC) surgery as previously described (Guo et al., 2021). Eyes, either right or left, were selected at random to receive experimental treatment, with unselected contralateral eyes used as controls. Briefly, for ONC, fine tweezers (Dumont#5; Fine Science Tools (FST) Item No. 11254-20 or Dumont#55; FST Item No. 11255-20) were used to create a small incision in the conjunctiva and then maneuvered between extraocular muscles to access the optic nerve, which was gently squeezed for 5 seconds at a location approximately 1mm posterior to the globe using Dumont #5 tweezers.

### 5.2. RGC Isolation

To isolate RGCs, we used a method that has previously been verified with slight modifications. To ensure enough tissue for proteomics, 5 retinas were pooled per experimental unit to create n=4 per group. Briefly, animals from each group were euthanized with CO_2_ asphyxiation and their retinas were harvested in cold neurobasal medium and placed in a 37°C water bath for 5 minutes. The neurobasal media was removed and replaced with fresh pre-warmed neurobasal media containing papain (0.06mg/ml or 33.4 U/mg) and 5mM L-cysteine and incubated 20 minutes at 37°C. The papain solution was removed and replaced with neurobasal containing 2mM L-Glutamine (Gibco; Catalog no. 25030081) B-27 supplement (Gibco; Catalog no. 17504044) and 10% FBS (Gibco; Catalog no. 26140079). The retinas were dissociated by gently pipetting up and down with a wide bore 1ml pipette tip. Cells were then centrifuged at 450g for 8 minutes and resuspended in 90 µl of isolation buffer (DPBS + 0.5% BSA + 2mM EDTA) containing 25 µg/ml DNAse I + 5mM MgCl2. Cells were filtered through a 30µm cell strainer and incubated with CD90.2 magnetic beads (Miltenyi Biotec; Catalog no. 130-121-278) at 4°C for 10 minutes and then isolated using MACs LS (Miltenyi Biotec; Catalog no. 130-042-401) columns following the manufacturer’s instructions. After isolation, cells were washed with DPBS, centrifuged at 450g for 8 minutes and their pellets stored at −70°C prior to proteomics sample preparation and LC/MS analysis.

### 5.3. Western Blotting

Enriched RGCs were lysed in RIPA buffer (Cell Signaling Technology, Cat: 9806) and protein concentrations were determined by Bio-Rad protein assay reagent (Cat: 5000006). Ten micrograms of protein were separated by SDS-PAGE and transferred to a PVDF membrane. After blocking with milk, membranes were then incubated with primary antibodies (1:2000 dilution) of interest (Rbpms-MilliporeSigma, Cat:ABN1362, Lot:3782213. TUJ1/TUBB3-Biolegend, Cat:801203, Lot:B373939. PDE6β-invitrogen, Cat:PA1-722, Lot: YA369623,; Rhodopsin-University of British Columbia, Rho 1D4, Lot: 1019. β-actin-MilliporeSigma, Cat: A2228, Lot:118M4829V), followed by HRP-conjugated secondary antibodies (Li-COR; Anti-mouse Cat: 926-80010, Lot:D20526-01 Anti-rabbit Cat: 926-80011, Lot: D10601-03) and lastly imaged on a Li-COR Odyssey Imaging System.

### 5.4. qRT-PCR

RNA was isolated from enriched RGCs using an RNeasy Micro Kit (Qiagen, 74004). RNA content was measured with a nanodrop spectrophotometer. cDNA was prepared using Bio-Rad iScript cDNA Synthesis Kit (1708891) following the manufacturer’s instructions. qRT-PCR was carried out on an Applied Biosystems QuantStudio 3. Predesigned primers were purchased from ThermoFisher (Gapdh-Mm99999915_g1, Rhodopsin-Mm01184405_m1 and Thy1-Mm00493681_m1).

### 5.5. Proteomics

#### 5.5.1. LC/MS

Proteomics analysis was carried out as previously described with minor changes (Ludwig et al., 2016), within section 2.5 nLC-ESI-MS2 under Protein IDs for GeLC. Proteins from isolated RGCs were extracted using T-PER™ Mammalian Protein Extraction Reagent (Thermo Fisher Scientific, Cat.# 78510) supplemented with HALT protease inhibitor cocktail (Thermo Fisher Scientific, Cat.# 78425), and benzonase nuclease (Sigma, E1014) following manufacturers instructions. Lysates were quantified using Pierce BCA Protein Assay Kit (Thermo Fisher Scientific, Cat.# 23227). Samples were prepared in NuPAGE LDS sample buffer (1x final conc., Invitrogen, Cat.# NP0007) and reduced with DTT then denatured at 70°C for 10min prior to loading 20µg onto Novex NuPAGE 10% Bis-Tris Protein gels (Invitrogen, Cat.# NP0315BOX) and separated appropriately (@ 200 constant V). The gels were stained overnight with Novex Colloidal Blue Staining kit (Invitrogen, Cat.# LC6025). Following de-staining, each entire lane was cut into multiple MW and equilibrated in 100 mM ammonium bicarbonate (AmBc), each gel plug was then digested overnight with Trypsin Gold, Mass Spectrometry Grade (Promega, Cat.# V5280) following manufacturer’s instruction. Peptide extracts were reconstituted in 0.1% Formic Acid/ddH2O at 0.1µg/µL.

Peptide digests (8µL each) were injected onto a 1260 Infinity nHPLC stack (Agilent Technologies), and separated using a 75 micron I.D. x 15 cm pulled tip C-18 column (Jupiter C-18 300 Å, 5 micron, Phenomenex). This system runs in-line with a Thermo Q Exactive HFx mass spectrometer, equipped with a Nanospray Flex™ ion source (Thermo Fisher Scientific), and all data were collected in CID mode. The nHPLC is configured with binary mobile phases that includes solvent A (0.1%FA in ddH2O), and solvent B (0.1%FA in 15% ddH2O/85% ACN), programmed as follows; 10min @ 5%B (2µL/min, load), 30min @ 5%-40%B (linear: 0.5nL/min, analyze), 5min @ 70%B (2µL/min, wash), 10min @ 0%B (2µL/min, equilibrate). Following each parent ion scan (300-1200m/z @ 60k resolution), fragmentation data (MS2) were collected on the top most intense 18 ions @7.5K resolution. For data dependent scans, charge state screening and dynamic exclusion were enabled with a repeat count of 2, repeat duration of 30s, and exclusion duration of 90s.

MS data conversion and searches The XCalibur RAW files were collected in profile mode, centroided and converted to MzXML using ReAdW v. 3.5.1. The mgf files were created using MzXML2Search (included in TPP v. 3.5) for all scans. The data was searched using SEQUEST (Thermo Fisher Scientific), which is set for three maximum missed cleavages, a precursor mass window of 20ppm, trypsin digestion, variable modification C @ 57.0293, and M @ 15.9949 as a base setting. Searches were performed with the mus musculus species specific subset of the UniProtKB database.

#### 5.5.2. Peptide filtering, grouping, and quantification

The list of peptide IDs generated based on SEQUEST search results were filtered using Scaffold (Protein Sciences, Portland Oregon). Scaffold filters and groups all peptides to generate and retain only high confidence IDs while also generating normalized spectral counts (N-SC’s) across all samples for the purpose of relative quantification. The filter cut-off values were set with minimum peptide length of >5 AA’s, with no MH+1 charge states, with peptide probabilities of >80% C.I., and with the number of peptides per protein ≥2. The protein probabilities will be set to a >99.0% C.I., and an FDR<1.0. Scaffold incorporates the two most common methods for statistical validation of large proteome datasets, the false discovery rate (FDR) and protein probability (Keller, Nesvizhskii, Weatherly). Relative quantification across experiments were then performed via spectral counting (Old, Liu), and when relevant, spectral count abundances will then be normalized between samples (Hyde).

#### 5.5.3. Statistical analysis

For generated proteomic data, two separate non-parametric-like statistical analyses were performed between each pair-wise comparison. These analyses included; A) the calculation of weight values by significance analysis of microarray (SAM; cut off >|0.8| combined with, B) T-Test (single tail, unequal variance, cut off of p < 0.05), which are then sorted according to the highest statistical relevance in each comparison. For SAM, the weight value (W) is a function statistically derived that approaches significance as the distance between the means (μ1-μ2) for each group increases, and the SD (δ1-δ2) decreases using the formula: W=(μ1-μ2)/(δ1-δ2). For protein abundance ratios determined with N-SC’s, we set a 1.5-fold change as the threshold for significance, determined empirically by analyzing the inner-quartile data from the control experiments using ln-ln plots, where the Pierson’s correlation coefficient (R) is 0.98, and >99% of the normalized intensities fell between the set fold change. Each of the tests (SAM, Ttest, and fold change) must be passed for to be considered significant.

## Author contributions

MG and CS designed the study. CS performed animal work. JM perfomed the proteomic study. MG, JM, and CS analyzed the data. MG and CS wrote the manuscript and prepared the figures.

